# Habenular μ-opioid receptor knockout and chronic systemic receptor blockade promote negative affect and heighten nociceptive sensitivity

**DOI:** 10.1101/2025.08.02.668171

**Authors:** EA Pekarskaya, JA Galiza-Soares, F Hough, CB Langreck, JM Tucciarone, JA Javitch

## Abstract

The μ-opioid receptor (MOR), a subtype of opioid G protein-coupled receptor, is expressed in multiple brain circuits and is particularly enriched in the habenula, a small epithalamic structure implicated in aversive states. MOR dysfunction has been linked to several psychiatric and nociceptive disorders. Identifying the key brain regions mediating the behavioral consequences of disrupted MOR signaling can shed light on the role of the opioid system in mood and pain regulation. In this study, we administered methocinnamox (MCAM), a long-acting, pseudo-irreversible MOR antagonist, acutely or chronically to adult C57BL/6J mice. A comprehensive behavioral battery was used to assess affective, social, and pain behavior. A single MCAM administration (10 mg/kg, s.c.) did not alter baseline behavior, but blocked opioid-induced analgesia, suggesting that basal µ-opioid tone does not contribute to these behaviors. In contrast, chronic MCAM administration (10 mg/kg, s.c., 3x/week for 4 weeks) led to increased anxiety-like behavior and decreased sociability, as well as enhanced mechanical allodynia and thermal hyperalgesia. Remarkably, selective knockout of habenular MORs in adult *Oprm1^fl/fl^* mice reproduced key features of the chronic MCAM phenotype, including anxiety-like behavior and mechanical hyperalgesia. Together, these findings reveal that sustained inhibition of MOR signaling disrupts affective and nociceptive processing and highlight the habenula as a node mediating key behavioral deficits of disrupted opioid signaling.

## INTRODUCTION

Opiates and synthetic opioids are widely used to treat pain conditions. However, the function of opioids extends beyond pain relief to modulating aspects of emotion and motivation. μ opioid receptors (MORs) are a subtype of inhibitory G protein-coupled receptors (GPCRs) activated by endogenous opioid peptides that have been shown to influence mood, sociability, and pain processing (1, 2). For example, some individuals with major depression exhibit altered MOR activation (3, 4). Chronic pain patients also show decreased MOR availability (5, 6) and the majority report comorbid depression and anxiety (7). Intriguingly, prolonged opioid use also causes alterations in opioid receptor expression, and withdrawal after discontinuation of opioids is associated with persistent negative affect and heightened pain sensitivity (8). A deeper understanding of how opioid signaling regulates baseline physiology and behavior is critical, as dysregulation of this system may underlie the development of maladaptive emotional and sensory states.

The consequences of MOR activation on brain function and behavior are dependent on the cell types expressing the receptor and their anatomical localization. Opioid receptors are enriched in brain circuits of pain, reward, stress, and affect (9, 10). Several reports have identified particularly dense MOR expression in the habenula (Hb) (9, 11, 12). Hb hyperactivity has been implicated in decreased motivation (13, 14), despair (15, 16), addiction (17, 18), stress (19, 20), and pain (21, 22). The Hb has two major subdivisions: medial and lateral (MHb and LHb, respectively). The MHb expresses high levels of MORs, and their activation reduces pain-related aversion (23). The LHb has become a region of interest in reward processing, and dysregulation of the area has been implicated in depression (16). The majority of studies investigating this region in depression to date have focused on LHb. Because most habenular MORs are located in the medial aspect (11), the specific contribution of Hb-MOR signaling to pain and affect remains underexplored.

Prior work has utilized pharmacological and genetic manipulations of MORs to identify behavioral phenotypes associated with the dysregulation of several opioid receptor subtypes. Constitutive MOR knockout (KO) mouse models have demonstrated significant behavioral changes, including i) increased exploratory and distressed behavior (24–26), ii) deficits in reciprocal social interaction (27), and iii) hyperalgesia to thermal stimuli and enhanced mechanical allodynia (25, 28). Given the potential confound of developmental compensation in these KO mice, non-selective MOR antagonists, such as naloxone and naltrexone, have also been used to explore the acute behavioral effects of opioid signaling disruption in adult mice (29, 30). However, their use complicates interpretation of behavioral results, given that δ- and κ-opioid receptors (DORs and KORs, respectively) are both antagonized by naloxone and naltrexone with known effects on behavior (31, 32). On the other hand, selective MOR antagonists, such as cyprodime, have other limitations, including lower potency and rapid metabolism, making these cumbersome for the study of chronic MOR disruption (33).

Methocinnamox (MCAM) is a pseudo-irreversible MOR antagonist that provides a long-lasting and highly selective *in vivo* blockade of MORs (34, 35). A single dose of MCAM induces MOR antagonism that lasts for at least three days and up to two weeks (34, 36). In the present study, we use MCAM to investigate the effect of acute versus chronic MOR blockade on affective, social, and nocifensive behaviors. A single administration of MCAM blocked opioid-induced analgesia without affecting other behaviors. In contrast, chronic MCAM treatment led to increased anxiety-like behavior and social deficits, as well as enhanced allodynia and hyperalgesia. To investigate the role of Hb MORs in maintaining these behaviors, we knocked down local Hb MORs in adulthood by injecting AAV-Cre-mCherry into the Hb of floxed MOR (*Oprm1^fl/fl^*) mice. Adult Hb-MOR knockout mice showed increased levels of anxiety-like behavior and mechanical hyperalgesia. These results suggest that maintaining basal tone at Hb-MORs is critical to the homeostasis of affective states and pain processing.

## METHODS

### Mice

Male C57BL/6J (Jackson Laboratory; Strain #:000664) and *Oprm1^fl/fl^* (Jackson Laboratory; Strain #:030074) mice were group-housed with ad libitum access to food and water (except during the novelty-suppressed feeding test). Animals were kept in a temperature and light-controlled environment with a 12/12 light/dark cycle. All mice (8-13 weeks old) were on a C57BL/6J background. Mice were purchased from Jackson Laboratories and housed in the New York State Psychiatric Institute or Stanford University School of Medicine animal facilities.

All procedures performed at Columbia were carried out per guidelines approved by the Institutional Animal Care and Use Committees at Columbia University and the New York State Psychiatric Institute. Procedures executed at Stanford were approved by Stanford University’s Administrative Panel on Laboratory Animal Care and the Administrative Panel of Biosafety.

### Drug preparation and administration

Methocinnamox (MCAM) (National Institute on Drug Abuse drug supply program, US; Tocris, UK) was dissolved in 10% w/v 2-hydroxypropyl-beta-cyclodextrin (Tokyo Chemical Industry, US; Merck, UK) vehicle (VEH) for administration. Either VEH or MCAM (10 mg/kg) was administered via subcutaneous (s.c.) injection once every three days to ensure sustained MOR blockade (37).

### Viral injections and stereotaxic surgery

AAVs were purchased from the Stanford Neuroscience Gene Vector and Virus Core: AAV-DJ-hSyn-Cre-eGFP, AAV-DJ-hSyn-eGFP, AAV-DJ-Ef1a-Cre-mCherry, and AAV-DJ-Ef1a-mCherry. Briefly, mice aged 7-9 weeks were anesthetized using isoflurane (1–2% v/v) and positioned in a stereotaxic apparatus (David Kopf Instruments, Tujunga, CA). Viral vectors were delivered bilaterally into the habenula (AP −1.55, ML ±0.25, DV −2.8 from Bregma) at a flow rate of 100 nL/min, with a total injection volume of 250 nL. Injections were carried out using borosilicate pipettes attached to a 5 μL Hamilton syringe mounted on a microinjection pump. Following a 5-minute diffusion period, pipettes were withdrawn slowly. Mice were allowed to recover for 3-4 weeks before behavioral testing.

### Immunohistochemistry

Mice were anesthetized using isoflurane and perfused transcardially with 1X PBS, followed by 4% paraformaldehyde in PBS (pH 7.4). The brains were extracted and post-fixed overnight in the same fixative. Subsequently, brains were sectioned at 50 μm using a vibratome, and the sections were collected in PBS. Free-floating sections were washed three times with PBS for 10 minutes at room temperature, then incubated for 1 hour in PBS containing 0.5% Triton X-100 and 5% normal goat serum. Following a 10-minute PBS wash, sections were incubated with primary antibodies—rabbit anti-MOR (1:300, abcam, ab134054) rabbit anti-TPH2 (1:1000, Novus Biologicals, NB100-74555), mouse anti-TH (1:1000, Millipore, MAB318), rat anti-mCherry (1:1000, Millipore, MAB131873), or chicken anti-GFP (1:1000, Aves Labs, GFP-1010)—in a carrier solution containing 0.5% Triton X-100 and 5% normal goat serum in PBS. This incubation was carried out with gentle shaking at room temperature for 24 hours. After four 10-minute washes in PBS, sections were incubated with species-specific secondary antibodies—Alexa Fluor 488, 594, or 647 (1:750, Invitrogen; A-11039, A-11058, A-31573)—in carrier solution for 2 hours at room temperature. Finally, sections were washed four times in PBS for 10 minutes each, mounted on SuperFrost Plus glass slides, and coverslipped using Fluoromount-G with DAPI (Southern Biotech, Birmingham, AL, 0100-20). Fluorescent images were acquired using a Nikon A1 confocal microscope or a Keyence BZ-X800 fluorescence microscope. Cells were counted and co-localization was measured using Fiji (38). The number of cells expressing MOR and mCherry was calculated within each Hb section and averaged for each subject, with three sections total per animal.

### Behavior

#### Open field

Animals were placed in a plexiglass open field chamber (42 x 42 x 38 cm) and allowed to explore for 60 minutes. Infrared arrays on the Y and X axes tracked animal movement, which was collected by the MotorMonitor software (Activity Monitor, Med Associates, Georgia, VT, US). Ambulatory movement, time spent in the center, and the number of rearing events were collected and analyzed. The center was defined as a central 21 x 21 cm grid. Center time and rearing events were summed over the first 10 minutes, while locomotion was measured over 60 minutes.

#### Marble burying

On testing day, each mouse was placed in a cage with 3 inches of corn cob bedding and 20 marbles lined up in four columns and five rows. Animals were allowed to explore for 30 minutes before being returned to their home cage. Two experimenters counted and confirmed the number of marbles buried, by at least 2/3rds, in the bedding.

#### Novelty-suppressed feeding

Mice were food deprived for a total of 15-18 hours before testing, as described previously (39). A pellet of food was placed on a white circular platform in the center of a bedding-covered arena (30 cm x 40 cm), with 1200 lux overhead lighting. The latency for each animal to enter the brightly lit center and bite the food pellet was recorded. After the first bite, the pellet was removed, and the animals remained in the apparatus for a total of 360 seconds. Immediately following the test, animals were placed individually into their home cages with access to a single pellet for five minutes. The latency to the first bite in the home cage and the total weight consumed were recorded to control for overall hunger during the arena task.

#### Automated Von-Frey

Cage mates were tested concomitantly, with each mouse placed in an individual enclosure (96mm x 96mm x 140mm) of the Dynamic Plantar Aesthesiometer (Ugo Basile, Gemonio VA, Italy). The device uses a filament to deliver a perpendicular force (1 - 10 g) at a chosen ramp time (1 s) to the plantar surface of the left hind paw. A response was considered positive if the animal exhibited any nocifensive behaviors in response, including brisk paw withdrawal, licking, or shaking of the paw.

#### Manual Von Frey

Cage mates were tested concomitantly, with each mouse placed in an individual custom-made pexiglass enclosure positioned on a homemade wire testing rack. Mechanical stimulation was applied to the plantar surface of the left hind paw using von Frey filaments (0.01 - 2 g plastic fibers, North Coast). A response was considered positive if the animal exhibited any nocifensive behaviors in response, including brisk paw withdrawal, licking, or shaking of the paw. Mechanical threshold values were calculated using the up-down method (40), which determines the 50% response threshold by oscillating stimulus intensity around the response level.

#### Hot plate

Each mouse was placed individually inside a Plexiglas enclosure on top of an aluminum plate (11" X 10.5" X ¾”, 275 mm X 263 mm X 15 mm) with a constant temperature of 55 + 0.2 °C (IITC Inc./Life Science Instruments, Woodland Hills, CA). The latency to flick or lick the hind paw was recorded with a maximum time limit of 30 seconds. The plate was wiped down with 70% ethanol between each animal.

#### Social interaction/novelty test

Animals were tested in a plexiglass open field chamber (42 x 42 x 38 cm). Mice were placed in the middle of the arena and allowed to explore for 5 minutes. Then, the animal was removed to a holding cage while two upside-down wire mesh pencil cups were placed on opposite sides of the arena. One pencil cup served as an empty container. The other pencil cup contained a novel 7-week-old C57BL/6J male mouse. Mice were then returned to the arena and allowed to explore for 5 minutes. Time spent exploring each cup and the total distance traveled were automatically quantified using ANY-maze software.

#### 3-chamber sociability test

The three-chamber sociability test was performed using a clear plexiglass apparatus divided into three interconnected chambers, as described previously (41). On the first day, experimental mice were habituated to the apparatus for 5 minutes with two empty wire mesh cups placed in the outer chambers. At the same time, age-, strain-, and sex-matched juvenile conspecifics (3–5 weeks old) were acclimated to the mesh cups for 5 minutes. On the second day, the experimental mouse was placed in the center chamber, and a juvenile conspecific was placed inside one of the wire cups. After a 2-minute acclimation period with the chamber doors closed, the barriers were raised, and the test mouse was allowed to freely explore all three chambers for a 20-minute session. Mouse location and movement were automatically recorded using BIOBSERVE video tracking software.

### Statistics

Data were analyzed using Prism (GraphPad, San Diego, CA). Data are expressed as mean ± S.E.M. except where otherwise stated. As appropriate, differences between groups were evaluated using a t-test (for continuous variables), a Mann-Whitney U-test (two-tailed) (for discrete variables), or a two-way ANOVA. The social interaction index was determined as follows: Social Interaction Index = (Time (novel mouse) – Time (empty cup))/(Time (novel mouse) + Time (empty cup)). The percent of maximal possible effect (%MPE) for tianeptine-induced analgesia in the hot plate was determined as follows: %MPE = (response − pre-value)/(cut-off time − pre- value) × 100.

## RESULTS

### Single-dose MCAM administration does not alter baseline behavior, but blocks opioid induced analgesia

To determine the effect of single-dose MCAM administration on pain, affective, and social behavior, C57BL/6J mice received injections of either 10 mg/kg MCAM or VEH on day 0. On day 1, animals were assessed in the open field (OF), social interaction (SI), and marble burying (MB) tests (Fig. 1A). Two weeks later, once the drug had completely cleared, the mice received another MCAM or VEH injection. The next day, the mice underwent pain behavioral testing with automated Von Frey (aVF) and hot plate (HP).

**Figure 1.**
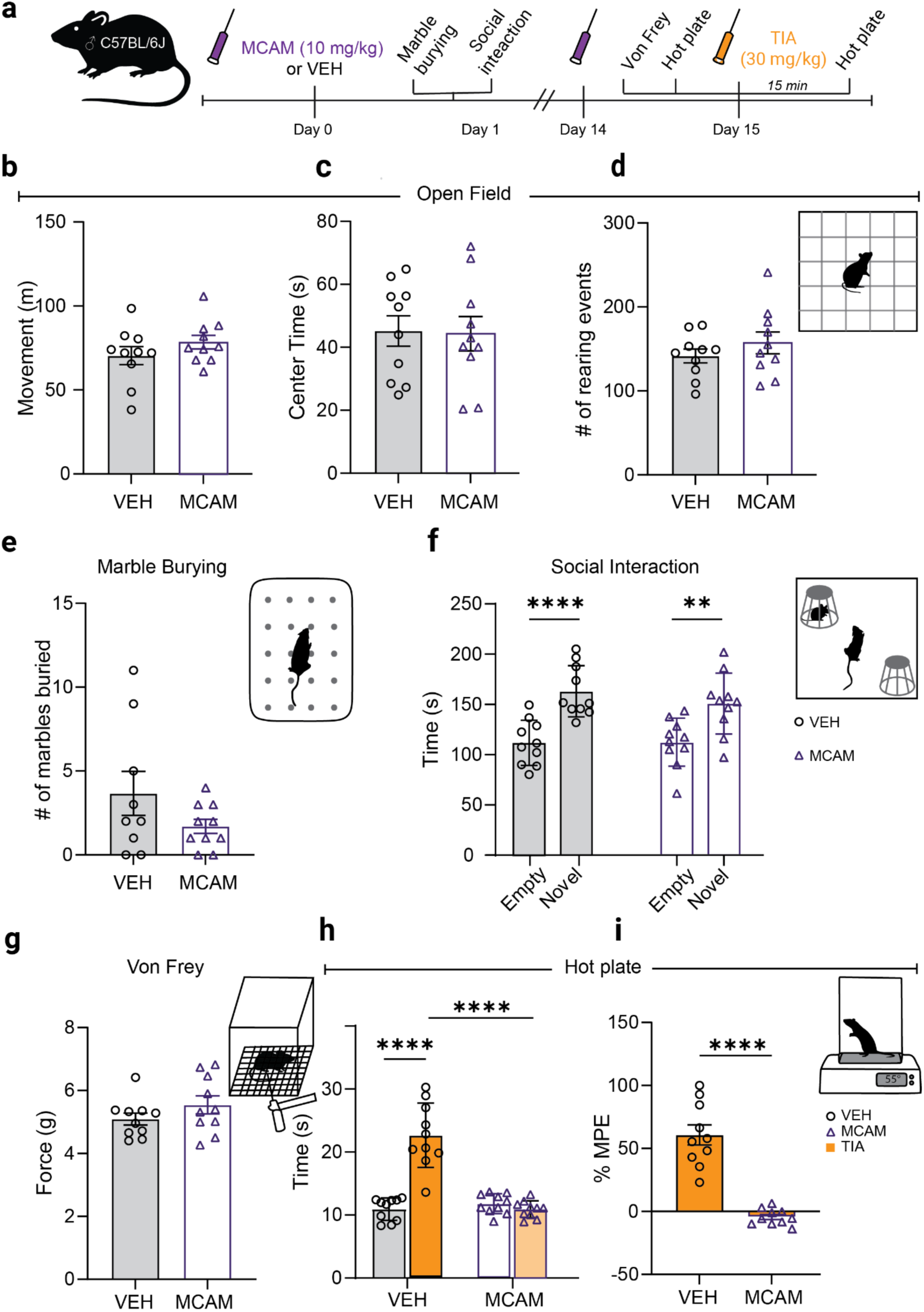
Acute MCAM administration does not alter baseline behavior but disrupts opioid- induced analgesia. a) Adult male C57BL/6J mice were subcutaneously injected with equal volumes of MCAM (10 mg/kg) or vehicle (s.c.). Subsequently, animals were subjected to behavioral testing. Mice were injected with tianeptine (30 mg/kg) after baseline hot plate to test opioid-induced analgesia. Subsequently, animals were subjected to behavioral testing. Behavioral battery included open-field, social interaction, marble burying, Von Frey, and hot plate. b) Total movement in 60 min of open-field (unpaired, two-tailed, t-test, p=0.2489, n=10 VEH, 10 MCAM). c) Center time in the first 10 min of open-field (unpaired, two-tailed, t-test, p=0.9050, n=10 VEH, 10 MCAM). d) Rearing events in the first 10 min of open-field (two-tailed, Mann-Whitney test, p=0.4467, n=10 VEH, 10 MCAM). e) Number of marbles buried (two-tailed, Mann-Whitney test, p=0.4047, n=9 VEH, 10 MCAM). f) Social interaction total time (two-way ANOVA, F (1, 36)=0.6307, p=0.4323 for interaction of treatment and social stimulus, F(1, 36)=30.29, ****p<0.0001 for main effect of social stimulus, F (1, 36)=0.5068, p=0.4811 for main effect of treatment, Šídák’s multiple comparisons post-hoc test for VEH: ***p=0.0002, for MCAM: **p=0.0040, n=10 VEH, 10 MCAM). Error bars represent standard error. *p<0.05, **p<0.01,***p <0.001, ****p<0.0001. g) Mechanical force threshold in Von Frey (unpaired, two-tailed, t-test, p=0.2095, n=10 VEH, 10 MCAM). h) HP latency to withdrawal, baseline vs acute tianeptine (two-way ANOVA, F (1, 18) = 64,84, ****p<0,0001 for interaction of analgesic administration and MCAM treatment, F (1, 18) = 49,79, ****p<0.0001 for main effect of analgesic administration, F (1, 18) = 26,27, ****p<0.0001 for main effect of MCAM treatment, Šídák’s multiple comparisons post-hoc test comparing baseline to tianeptine treatment for VEH: ****p<0.0001, for MCAM: ****p<0.7400; n=10 VEH, 10 MCAM). i) HP MPE % (unpaired, two-tailed, t-test, ****p=0.<0001, n=10 VEH, 10 MCAM). Error bars represent standard error. *p<0.05, **p<0.01,***p <0.001, ****p<0.0001.

The OF test was used to assess anxiety-like behavior and overall locomotion. There was no difference between the VEH and MCAM groups in overall locomotion (Fig. 1B, t-test, p=0.2489), center time (Fig. 1C, t-test, p=0.9050), or rearing events (Fig. 1D, Mann-Whitney test, p=0.4467). Single-dose administration of MCAM also did not significantly alter the number of marbles buried, which is often used as a measure of anxiety or perseverative behavior (42) (Fig. 2E, Mann-Whitney test, p=0.4047).

**Figure 2.**
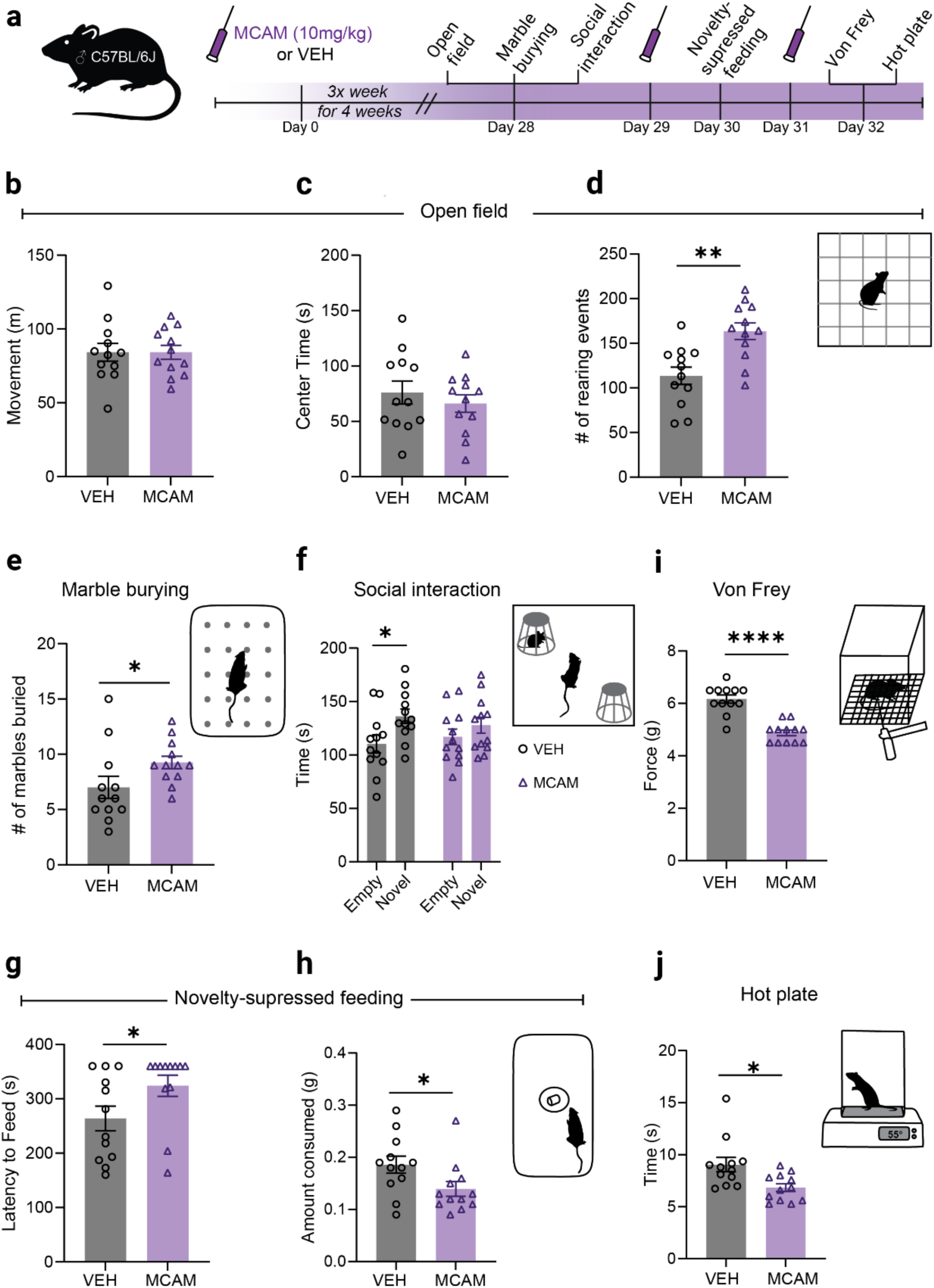
Chronic MCAM administration alters anxiety-like, social, and pain behaviors. (a) Equal volumes of MCAM (10 mg/kg) or vehicle were subcutaneously (s.c.) injected every 3 days for 4 weeks in adult male C57BL/6J mice. Subsequently, animals were subjected to behavioral testing. Behavioral battery included open-field, social interaction, marble burying, Von Frey, and hot plate. (b) Total movement in 60 min of open-field (unpaired, two-tailed, t-test, p=0.9991, n=12 VEH, 12 MCAM). (c) Center time in the first 10 min of open-field (unpaired, two-tailed, t-test, p=0.4516, n=12 VEH, 12 MCAM). (d) Rearing events in first 10 min of open-field (two-tailed, Mann Whitney test, **p=0.0022, n=12 VEH, 12 MCAM). (e) Number of marbles buried (two-tailed, Mann Whitney test, p=0.0198, n=12 VEH, 12 MCAM). (f) Social interaction total time (two-way ANOVA, F (1, 44) = 0.9957, p=0.3238 for interaction of chronic treatment and social stimulus, F (1, 44) = 6.041, *p=0.0180 for main effect of social stimulus, F (1, 44) = 0.01341, p=0.9083 for main effect of treatment, Šídák’s multiple comparisons post-hoc test for comparing time spent with the empty cup vs. social stimulus VEH: *p=0.0369, for MCAM: 0.5205, n=12 VEH, 12 MCAM). (g) Latency to feed in the novelty-suppressed feeding task (two-tailed, Mann Whitney test, p = 0.0473, n=12 VEH, 12 MCAM). (h) Home cage consumption in NSF (unpaired, two-tailed, t-test, *p=0.0416, n=12 VEH, n=12 MCAM). Error bars represent standard error. *p<0.05, **p<0.01,***p <0.001, ****p<0.0001. (i) Mechanical Von Frey threshold (unpaired, two-tailed, t-test, ****p<0.0001, n=12 VEH, 12 MCAM). (j) Hot plate latency to withdrawal (unpaired, two-tailed, t-test, p=0.0105, n=12 VEH, 12 MCAM). Error bars represent standard error. *p<0.05, **p<0.01,***p <0.001, ****p<0.0001.

Because MOR activity has been implicated in social behavior (43), we next tested the effect of a single-dose MCAM on sociability using the SI test. There was no difference between VEH and single-dose MCAM administration in terms of time spent with a novel conspecific vs. an empty cup (Fig. 1F, two-way ANOVA, p=0.4323). Both the MCAM and VEH groups showed a clear preference for the social stimulus, with both spending more time near the novel mouse. In contrast, non-specific opioid antagonists, such as naloxone, have been shown to impact sociability when administered acutely (44), presumably through its actions at other opioid receptors.

A single dose of MCAM has been previously shown to block morphine-induced analgesia without altering baseline mechanical withdrawal latency in rats (45). We next aimed to replicate these findings in mice, utilizing the atypical antidepressant tianeptine (TIA), which is a MOR partial agonist shown to produce MOR-dependent analgesia (46, 47). On day 14, after a two-week washout period, the same cohort received a second MCAM injection. Twenty-four hours later, animals underwent the aVF test, followed by HP testing. Single-dose MCAM administration did not change the mechanical pain threshold in the aVF test compared to the VEH group (Fig. 1G, t-test, p=0.2095), nor did it alter baseline HP measurements. Next, animals were administered TIA (30 mg/kg, i.p.) and then assessed for response latency in the HP after 15 minutes. We observed a significant difference in analgesic response between the VEH and MCAM-treated groups (Fig. 1I, t-test, p=0.<0001), with the VEH-treated animals showing significant TIA-induced analgesia and MCAM treatment resulting in no significant change in response from baseline (Fig. 1H, two-way ANOVA, p<0.0001).

Overall, MOR-blockade through a single dose of MCAM did not generate negative affective behavioral states or alter baseline nociception. However, it blocked opioid-induced analgesia 24 hours after administration, confirming MCAM’s long-lasting action as an MOR antagonist.

### Chronic MCAM administration alters affect, social behavior, and nociception

Next, we sought to determine the effects of sustained selective MOR blockade on pain, affective, and social behavior. C57BL/6J mice received injections of either 10 mg/kg MCAM (n = 12) or VEH (n = 12) every 3 days for 4 weeks, an interval we have previously demonstrated to be effective for sustained functional MOR blockade (37). After 4 weeks, animals were subjected to a comprehensive behavioral battery (Fig. 2A). The following behavioral tests were performed, with each test being given at least 24 hours after the last MCAM injection: OF, MB, SI, novelty-suppressed feeding (NSF), aVF, and HP tests.

In the OF test, there was no difference in the overall distance traveled in 60 minutes between the VEH and MCAM groups (Fig. 2B, t-test, p=0.9991), suggesting that chronic MCAM does not alter overall locomotor ability. We also did not observe a difference in center time between the groups (Fig. 2C, t-test, p=0.4516). There was, however, a significant difference in the number of rearing events, with MCAM-treated mice showing increased rearing as compared to controls, indicating increased anxiety-like and/or exploratory behavior (48) (Fig. 2D, Mann Whitney test, p=0.0022). Additionally, MCAM-treated mice buried significantly more marbles than VEH-treated mice, suggesting that chronic MCAM treatment increases anxiety-like and/or perseverative behavior (Fig. 2E, Mann Whitney test, p=0.0198).

While there was no significant interaction between the treatment group and the object in the SI test (Fig. 2F, two-way ANOVA, F (1, 44) = 0.9957, p=0.3238 for interaction of chronic treatment and social stimulus), MCAM mice spent equal amounts of time near the novel mouse and the empty cup (Fig. 2F, MCAM: Šídák’s multiple comparisons test, p = 0.5205). In contrast, VEH-treated mice showed a clear preference for the novel conspecific mouse (Fig. 2F, VEH: Šídák’s multiple comparisons test, p = 0.0369), indicating that sustained MOR blockade impaired sociability.

As an additional measure of anxiety-like behavior, the animals were tested in the NSF test. To dissociate anxiety-like behavior from appetite or metabolic changes, we also measured home cage feeding to assess latency to eat and total food consumption in a familiar, low-anxiety environment as a control. We found that latency to feed in the NSF test was significantly higher in the chronic MCAM group (Fig. 2G, Mann Whitney test, p = 0.0473). However, we also observed an overall decrease in home-cage consumption (Fig. 4H, t-test, p=0.0416), making it difficult to discern if the latency to first bite in the NSF test is due to decreased hunger or increased anxiety-like behavior.

Finally, we assessed whether repeated MCAM administration would impact baseline nocifensive responses. On day 31, mice were subjected to the aVF and HP tests to assess mechanical and thermal pain responses, respectively. Remarkably, MCAM-treated mice showed a steep reduction in mechanical pain threshold in the aVF test (Fig. 2I, t-test, p<0.0001). VEH-treated mice required an average of 6.17 g to show a nocifensive reaction, while MCAM-treated mice only needed an average force of 4.89 g. The MCAM group also showed decreased latency to jump or lick/shake their hind paws in the HP test compared to controls (Fig. 2J, t-test, p=0.0105). Thus, it appears that chronic MOR blockade can impair baseline pain regulation, leading to heightened mechanical and thermal sensitivity.

### MORs are densely expressed in the habenula and can be targeted via a viral approach

Given that chronic systemic MOR blockade produced significant changes in affect and nociception, we aimed to identify a candidate brain region where disrupted MOR signaling would similarly influence behavior. The Hb emerged as a strong candidate due to its well-established role in motivation and pain processing (49). Additionally, this brain region is reported to express high levels of MORs (11), though quantitative data on receptor distribution within the Hb remain limited. To address this, we used immunohistochemistry to quantify MOR expression across the anterior-posterior (A-P) axis of the Hb. Consistent with prior reports, we observed dense MOR labeling, particularly in the medial aspect and at the border between MHb and LHb (Fig. 3A). MORs were present throughout the full A-P extent of this brain region, with higher densities around the −1.55 and −1.79 coordinates (Fig. 3B).

**Figure 3.**
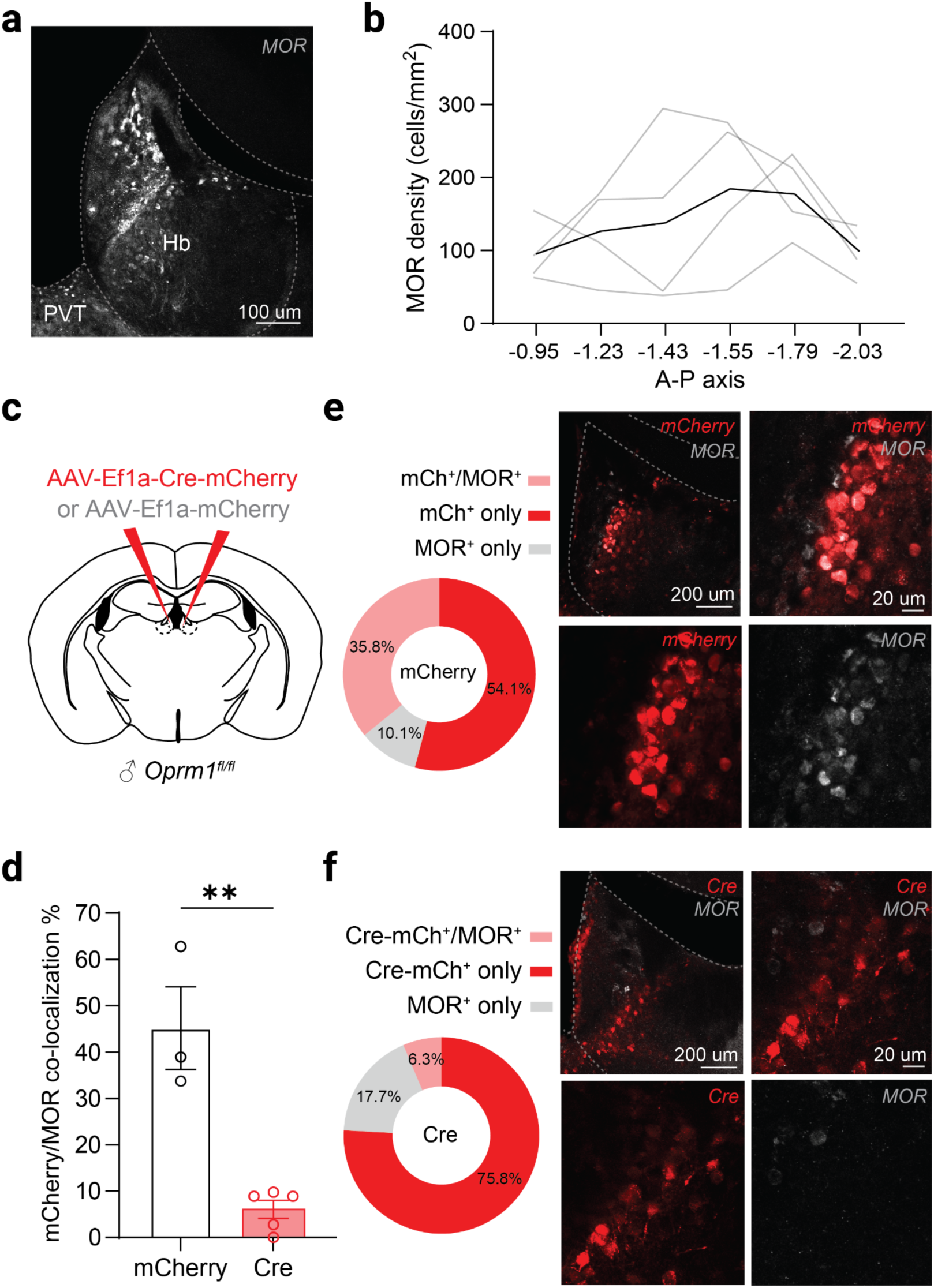
MOR expression and targeted *Oprm1* deletion in the habenula. (a) Representative coronal brain section of the habenula (A-P coordinate = −1.55) showing MOR immunoreactivity. (b) Quantification of MOR density per mm^2^ across the anterior-posterior axis. (n = 4). (c) Schematic representation of viral injection sites in *Oprm1^fl/fl^* mice, using either Cre or control virus. (d) % colocalization of mCherry reporter and MOR in control and Cre-expressing mice (unpaired, two-tailed, t-test, **p=0.0014, n=3 mCherry, 5 Cre). (e) Left, representative coronal brain section showing co-localization of MOR immunofluorescence with mCherry reporter expression. Right, pie chart showing high degree of mCherry/MOR colocalization in control mice (n = 477). (f) Left, representative coronal brain section demonstrating Cre-dependent *Oprm1* deletion, indicated by reduced MOR immunoreactivity in mCherry+ cells. Right, pie chart showing low degree of mCherry/MOR colocalization in Cre-expressing mice (n = 1259). Error bars represent standard error. *p<0.05, **p<0.01,***p <0.001, ****p<0.0001.

To selectively delete *Oprm1* in MOR-expressing cells within the Hb, we sought a viral vector capable of effectively infecting this region. As previously reported, the Hb can be resistant to many commonly used viral serotypes and promoters (50), possibly due to its remarkable cellular heterogeneity. In particular, the MHb has been shown to be resistant to most serotypes (51), prompting us to utilize AAV-DJ, a hybrid vector of eight AAV serotypes. We initially tested AAV-DJ vectors carrying either hSyn-Cre-eGFP or hSyn-eGFP to induce μ-opioid receptor deletion (Supplementary Figure 2A). However, both constructs yielded negligible reporter expression in the MHb, with transduction largely restricted to cells in the LHb. This pattern resulted in minimal overlap with MOR-expressing cells (Supplementary Figure 2D), suggesting the need for a promoter with broader efficacy across Hb subregions. We therefore turned to the elongation factor 1 alpha (Ef1a) promoter, which is robustly expressed throughout both the MHb and LHb (52, 53).

To selectively delete MORs, we injected AAV-DJ-Ef1a-Cre-mCherry or AAV-DJ-Ef1a-mCherry into the Hb of *Oprm1^fl/fl^* mice (Fig. 3C). This approach yielded robust reporter expression across the Hb. To assess the efficiency of receptor deletion, we quantified mCherry and MOR co-expression at the cellular level. Injection of the Cre virus significantly decreased the mCherry/MOR co-localization when compared to controls (Fig. 3D). Specifically, Cre-expressing mice showed markedly reduced MOR immunoreactivity in mCherry+ cells compared to the control group, with around 45.2% ± 8.9% overlap in controls and only 6.1% ± 4.4% overlap in the Cre group. As expected, MOR+ cells were overall present to a much lower degree in the Cre group (Fig. 3E-F). These results confirm both the abundance of MOR expression in the Hb and the effectiveness of our viral strategy in conditionally deleting MORs in adult mice.

### Deletion of habenular MORs in adulthood alters affect and nociception

To determine whether deletion of habenular MORs would impact pain, affective, and social behavior, mice were subjected to a behavioral battery focusing on aspects that were disrupted with chronic systemic pharmacological blockade, including MB, SI, VF, and HP (Fig. 4A-B).

**Figure 4.**
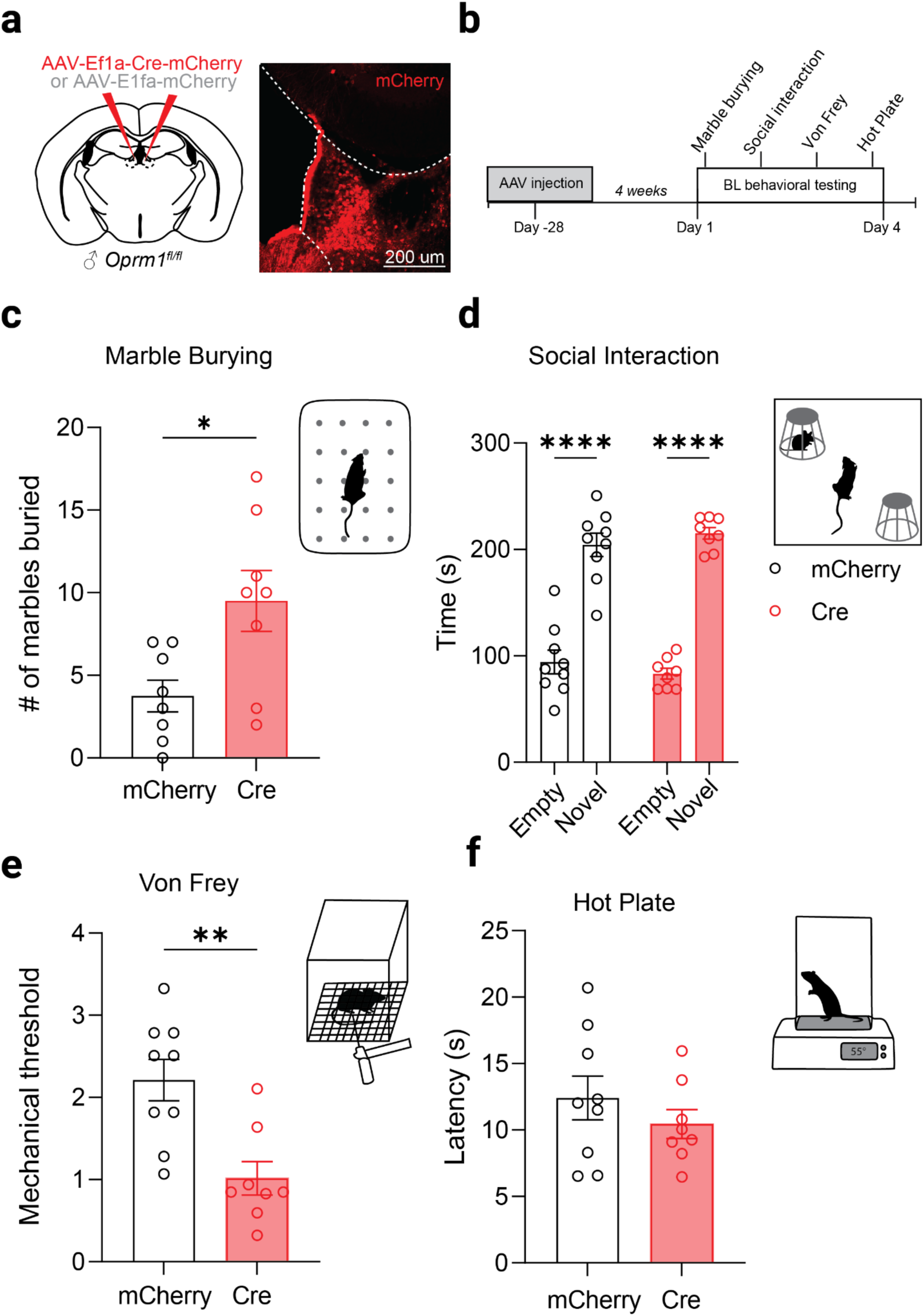
Adult *Oprm1* knockout in the habenula alters anxiety-like behavior and mechanical allodynia. (a) Coronal schematic and representative histological image showing the injection site in the habenula of *Oprm1^fl/fl^* male mice, following delivery of Cre or control virus. (b) Tests used to assess affective, social, and pain behavior. Behavioral battery included marble burying, social interaction, manual Von Frey, and hot plate. (c) Number of marbles buried (two-tailed, Mann Whitney test, *p=0.0190, n=9 mCherry, 8 Cre). (d) Social interaction total time (two-way ANOVA, F (1, 30) = 1.482, p=0.2330 for interaction of knockout and social stimulus, F (1, 30) = 180.2, ****p<0.0001. for main effect of social stimulus, F (1, 30) < 0.0001, p=0.9976 for main effect of knockout). Error bars represent standard error. *p<0.05, **p<0.01,***p <0.001, ****p<0.0001. (e) Mechanical Von Frey threshold (unpaired, two-tailed, t-test, **p<0.001, n=9 mCherry, 8 Cre). (f) Hot plate latency to withdrawal (unpaired, two-tailed, t-test, p=0.3512, n=9 mCherry, 8 Cre). Error bars represent standard error. *p<0.05, **p<0.01,***p <0.001, ****p<0.0001.

Mice expressing the active Cre buried significantly more marbles than the control mCherry group in the MB test (Fig. 4C, Mann Whitney test, p=0.019). Specifically, Cre-expressing mice buried an average of 9 ± 2 marbles, a similar increase to mice chronically treated with MCAM (Fig. 2E), suggesting that MOR signaling in the Hb contributes to the regulation of anxiety-like and/or perseverative behaviors.

Hb-MORs have been previously reported to contribute to social reward (54), however, whether deletion of MORs in this brain region during adulthood can impair sociability was not clear. To evaluate the impact of Hb-MOR deletion on sociability, we first performed a standard SI assay. Both the Cre and control groups demonstrated a clear preference for interacting with a novel conspecific over an empty cup (Fig. 4D, two-way ANOVA, p<0.0001). These data indicate that deletion of MORs in the Hb does not impair baseline sociability. To validate this finding, mice were also assessed using the three-chamber sociability (3CS) test (Supplementary Figure 3), which further confirmed intact social preference (Supplementary Figure 3B, two-way ANOVA, p<0.0001). Thus, Hb-MORs are not required for normal social approach behavior in adult mice.

We also assessed whether MOR deletion in the Hb would affect nociceptive processing. Although a role for the Hb in pain modulation has been described across species (55), the exact contribution of MOR signaling within this region remains poorly defined. Notably, Hb-MOR KO mice displayed significantly reduced mechanical allodynia thresholds relative to controls (Fig. 4E, t-test, p < 0.001), indicating that endogenous MOR activity in the Hb contributes to baseline mechanical nociception. Further dissection of subregional involvement revealed that deletion of MORs confined to the LHb did not affect mechanical sensitivity (Supplementary Figure 2C, t-test, p = 0.8796), suggesting a critical role for the MHb in this process. Thermal pain responses were assessed via HP test. No significant differences were observed in hind paw withdrawal latency between Cre and control groups (Fig. 4F, t-test, p = 0.3512), indicating that Hb-MORs may selectively modulate mechanical, but not thermal, nociceptive pathways.

## Discussion

Clinical and preclinical data suggest that reduced MOR activation/availability is associated with negative emotional states, including those associated with psychiatric disorders such as major depression and anxiety (4, 43, 56), as well as chronic pain conditions, such as neuropathic pain and fibromyalgia (5, 6, 57). In this study, we investigated how the duration and anatomical specificity of MOR signaling disruption influence affective, social, and nociceptive behaviors in adult C57 mice. Our findings demonstrate that a single systemic dose of the long-acting MOR antagonist MCAM does not significantly alter affect and pain-associated behaviors, whereas chronic systemic MOR blockade induces increased anxiogenic behavior, social avoidance, and sensitivity to both mechanical and thermal stimuli. Furthermore, selective knockout of MORs in the medial habenula is sufficient to recapitulate several aspects of chronic systemic MOR blockade, including increased anxiogenic behaviors such as repetitive digging and increased sensitivity to mechanical stimuli. These results suggest that ongoing MOR signaling is critical for maintaining emotional and sensory homeostasis and that chronic disruption of MOR function, particularly when localized to the medial habenula, may promote maladaptive phenotypes relevant to chronic pain and mood disorders.

The strength of these findings lies in the use of both systemic pharmacological and spatially restricted genetic approaches to dissect the role of MOR signaling in adult behavior. Pharmacological MOR blockade and constitutive *Oprm1* KO models have provided valuable insights into both the immediate and long-term role of endogenous opioids on behavior. However, most commonly used antagonists, such as naloxone and naltrexone, lack MOR selectivity and bind to δ-and κ-opioid receptors, complicating interpretation due to the known behavioral effects of these other receptor subtypes. Other MOR antagonists—such as cyprodime, beta-funaltrexamine, and CTAP—are primarily used for ex vivo or acute studies, but they are limited by rapid metabolism and/or insufficient selectivity (33, 58, 59), making them unsuitable for investigating chronic MOR blockade. Unlike conventional antagonists, MCAM offers greater selectivity for MORs and a pseudo-irreversible mechanism of action, enabling chronic antagonism with less frequent dosing. While MCAM blocks μ, δ, and κ agonist-induced analgesia an hour after administration, by 24 hours post-administration, it selectively inhibits MOR-mediated effects (60, 61). To minimize off-target effects and enhance interpretability, all behavioral tests were conducted 24 hours post-injection. Nonetheless, the potential impact of repeated transient κ and δ receptor antagonism warrants further study. We found that chronic pharmacological blockade of MORs led to a recapitulation of some but not all of the previously observed behavioral changes in global Oprm1^-/-^ mice. Similar to Oprm1^-/-^ mice, we show that chronic MCAM mice bury more marbles, spend less time around novel conspecifics, and have a higher latency to feed in the NSF test (62). Chronic MCAM mice show decreased consumption in the home cage. Unlike previous work showing increased time spent in the open arms of the elevated plus maze, chronic MCAM mice did not spend more time in the center of the OF (two metrics that often approximate the same exploratory and anxiety-like behavior in rodents) (24). Chronic MCAM mice also exhibited a significant and consistent increase in rearing behavior, a complex behavior that can indicate either heightened anxiety or increased exploratory drive. In anxiety-related contexts, it may reflect elevated vigilance or stress (48). Finally, we found that chronic MOR blockade resulted in hyperalgesia and enhanced allodynia, consistent with what is seen in some Oprm1^-/-^ mice (25, 28, 63), but not all (64). Taken together, these findings demonstrate that chronic disruption of μ-opioid receptor signaling can produce lasting alterations in mood, social behavior, and pain sensitivity—domains commonly affected in chronic pain conditions, mood disorders, and OUD.

From a translational perspective, these findings are particularly salient given renewed interest in MOR antagonists as novel OUD treatments. MCAM has recently emerged as a promising candidate for OUD treatment due to its long duration of action and MOR specificity. Chronic MCAM administration has reduced fentanyl self-administration in rhesus monkeys without significantly affecting feeding, heart rate, blood pressure, or body temperature (65). However, our findings suggest that long-term MCAM use may impact affective, social, and nocifensive behaviors, highlighting the need for further safety and side effect evaluation. In clinical contexts, these side effects, such as increased anxiety, social withdrawal, or heightened pain sensitivity, could undermine treatment adherence and overall outcomes, particularly during long-term maintenance therapy. Moreover, our data indicate that endogenous MOR signaling is critical for the maintenance of baseline mood and sensory function even in the absence of exogenous opioids, highlighting the need for careful evaluation of long-acting antagonists in vulnerable patient populations.

Additionally, the use of adult-onset, region-specific MOR knockout provides a novel avenue for evaluating MOR function while avoiding developmental compensations that confound constitutive Oprm1 knockout models, limiting their relevance to conditions that manifest in adulthood, such as chronic pain and opioid use disorder. To address these limitations and investigate the behavioral consequences of disrupted μ-opioid signaling in the mature brain, we used an adult-onset conditional MOR KO model. We also selected the Hb as a candidate brain region in which disruption of MOR signaling might lead to deficits in affective, social, and pain-related behaviors, as the region is widely recognized as a key node in the regulation of aversive states (66). However, the specific contributions of MOR-expressing neurons within this region remain poorly understood. Given the high density of MORs in the Hb and its proposed involvement in nociceptive and affective processing (55), investigating this population offers a valuable opportunity to elucidate how disrupted MOR signaling may contribute to mood disturbances, social dysfunction, and altered pain sensitivity.

A key finding of this study is that deletion of Hb-MORs is sufficient to alter baseline nociception, even in the absence of a pathological pain state such as neuropathic injury. Our data contribute to growing evidence implicating the habenula as a hub for integrating affective and nociceptive states. Consistent with prior work, we found that MORs are densely expressed along the border between the medial habenula (MHb) and lateral habenula (LHb) (11), and their deletion elevated rearing and marble burying behavior, phenotypes often associated with anxiety and compulsivity. These behavioral changes are consistent with a model in which MORs exert tonic inhibitory control over excitatory habenular neurons. Given that the habenula is predominantly glutamatergic, and that MORs are Gi/o-coupled receptors, loss of inhibitory MOR signaling likely enhances habenular output. Indeed, recent evidence shows that increased excitatory drive from the Hb contributes to both depression- and pain-like behaviors, and that chemogenetic silencing of this output can alleviate such phenotypes (21).

Notably, the behavioral consequences of chronic MCAM only partially overlapped with those observed following genetic MOR knockout in the Hb. Both models produced anxiogenic phenotypes and mechanical hypersensitivity, suggesting that disruption of endogenous MOR signaling in this region is sufficient to drive key affective and nociceptive impairments. In contrast, social behavior deficits were only observed with systemic MCAM administration, not with habenular MOR deletion. This suggests that social deficits may arise either from MOR disruption in other brain regions (e.g., nucleus accumbens or amygdala) or through broader network-level effects of systemic MOR blockade. It is also possible that MORs in the Hb modulate social behavior primarily under conditions of stress or withdrawal. Supporting this, MOR deletion in GPR151-expressing Hb neurons was recently shown to impair social preference only after social defeat stress (49).

More recently, an indirect pathway connecting glutamatergic Hb neurons to dopaminergic neurons in the VTA has been shown to mediate affective and nociceptive deficits in neuropathic pain models (21). However, most of this work has focused on the LHb, while the MHb, where MOR expression is most enriched, remains largely overlooked. To our knowledge, this is the first study to demonstrate a role for Hb MORs in maintaining basal nociceptive threshold. Notably, MOR deletion in the LHb did not produce similar effects on pain response, suggesting that previously reported pain-related functions of the LHb may involve distinct non-opioidergic mechanisms or different neuronal populations. The DRN has been implicated in pain processing (67), while the IPN has been mainly implicated in mood regulation (66). It is possible MOR- expressing neurons in the Hb may contribute to DRN-mediated nociceptive circuits. Additional work is necessary to investigate the brain circuits mediating Hb’s effect on nociception, as well as how it may be exacerbated in chronic pain states.

Our findings also complement recent work by Margolis and colleagues demonstrating that lateral preoptic area (LPO) projections to the habenula are modulated by presynaptic MORs, suggesting that terminal MORs may regulate excitatory input into this structure (68). In contrast, our approach targeted postsynaptic MORs within the habenula itself, enabling us to isolate the intrinsic contributions of local MOR signaling to affective and sensory behavior. The combined evidence suggests a dual modulatory role for MORs in this circuit: presynaptic inhibition of excitatory LPO input, and postsynaptic regulation of habenular excitability. Notably, MCAM would target both populations of MOR, while our knockout strategy was limited to postsynaptic MOR. Future studies using projection-specific manipulations of MORs will be critical to disentangle these convergent mechanisms. Moreover, our findings contrast with the known behavioral effects of tianeptine, an atypical antidepressant that acts as a MOR agonist. Tianeptine has been shown to alleviate pain in spared nerve injury models (69), improve depressive-like behaviors in rodents, and demonstrate efficacy in patients with major depressive disorder. Unpublished work further suggests it ameliorates the affective consequences of neuropathic pain (70). Given that tianeptine enhances MOR signaling, its behavioral effects may reflect the inverse of the phenotypes observed following MCAM treatment or MOR knockout, supporting a model in which MOR tone in the habenula acts as a critical regulator of pain and mood homeostasis. Moreover, we recently showed that anti-stress effects of (*R,S*)-ketamine and the more selective N-methyl-D-aspartate receptor (NMDAR) antagonist fluoroethylnormemantine (FENM) are also dependent on MOR signaling (37).

In conclusion, our results underscore the essential role of ongoing MOR signaling in maintaining emotional and nociceptive balance in adulthood. Acute MOR blockade is well tolerated, but chronic disruption—whether systemic or localized—induces behavioral alterations reminiscent of those observed in chronic pain and mood disorders. These findings deepen our mechanistic understanding of endogenous opioid function and highlight both opportunities and potential caveats in the therapeutic use of long-acting MOR antagonists. Future work should expand on the circuit-level substrates of these effects and evaluate the behavioral consequences of MOR disruption in disease-relevant contexts, such as injury, stress, or opioid withdrawal.

## Acknowledgements

This work was supported by the Hope for Depression Research Foundation (J.A.J.), by NIH grant F31 NS127547 (E.A.P.), and by NIH grant K08 DA055157, a Brain and Behavior Research Foundation Young Investigator Grant, The William and Katharine Duhamel Addiction Medicine Fund, and the Stanford University School of Medicine Department of Psychiatry & Behavioral Sciences 2024 Innovator Grants Program (J.M.T).

## Figures

**Supplementary Figure 1.**
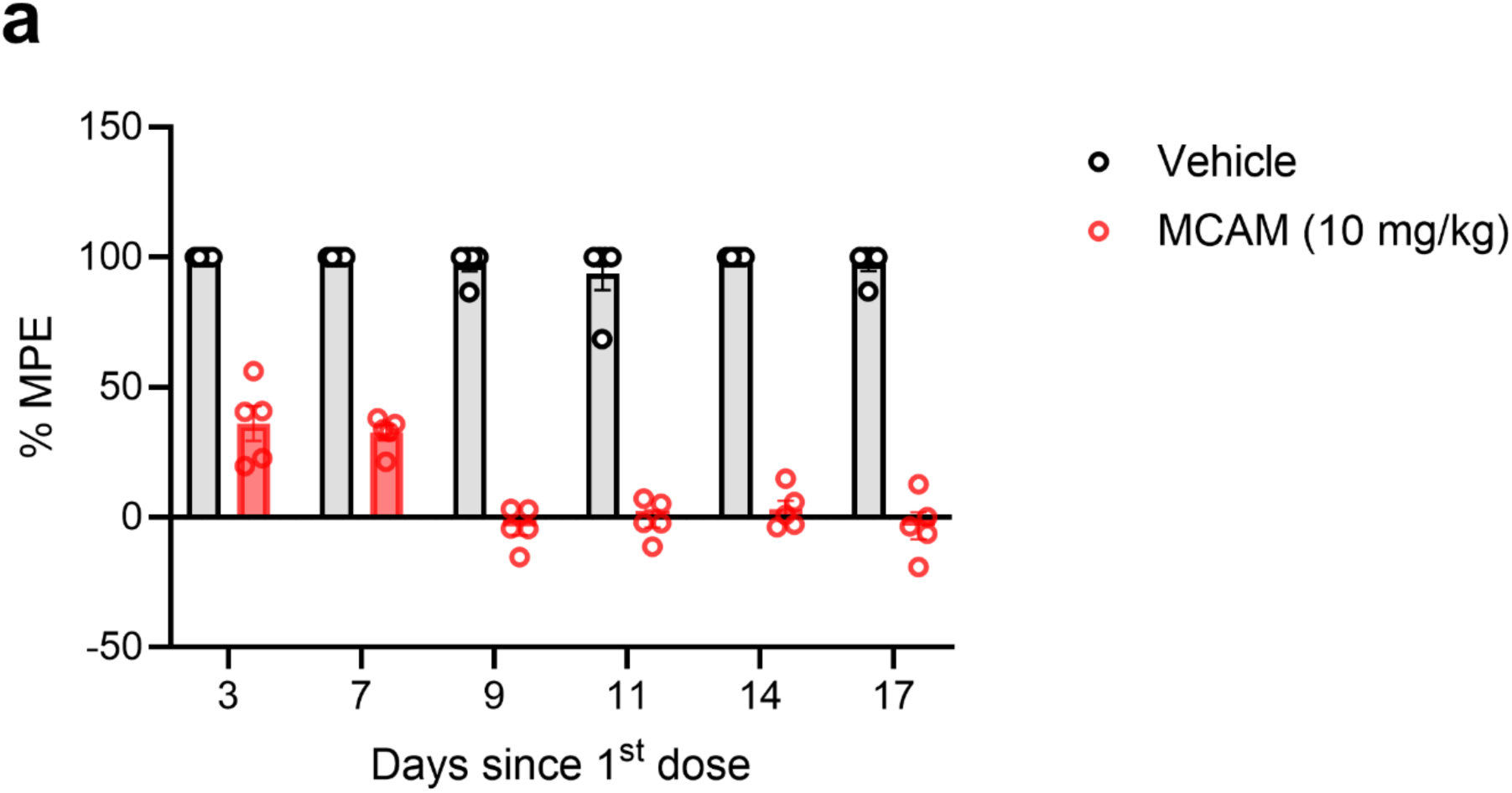
MCAM dosing pilot. Mice were administered MCAM (10 mg/kg, i.p.) or vehicle 24 hours before testing response latency in the hot plate following acute opioid administration. (a) Hot plate latency in response to morphine (30 mg/kg i.p.) with repeat MCAM administration every 2-4 days for 17 days. Mice were tested on the indicated days before being redosed with MCAM to assess sustained MOR blockade (two-way ANOVA, F (5, 40) = 9.581, ****p<0.0001 for interaction of days post first injection and treatment group, F (2.7, 21.6) = 13.46, ****p<0.0001 for main effect of days post 1st injection, F (1, 8) = 2716, ****p<0.0001 for treatment group, n = 5 vehicle, 5 MCAM). Error bars represent standard error. *p<0.05, **p<0.01,***p <0.001, ****p<0.0001.

**Supplementary Figure 2.**
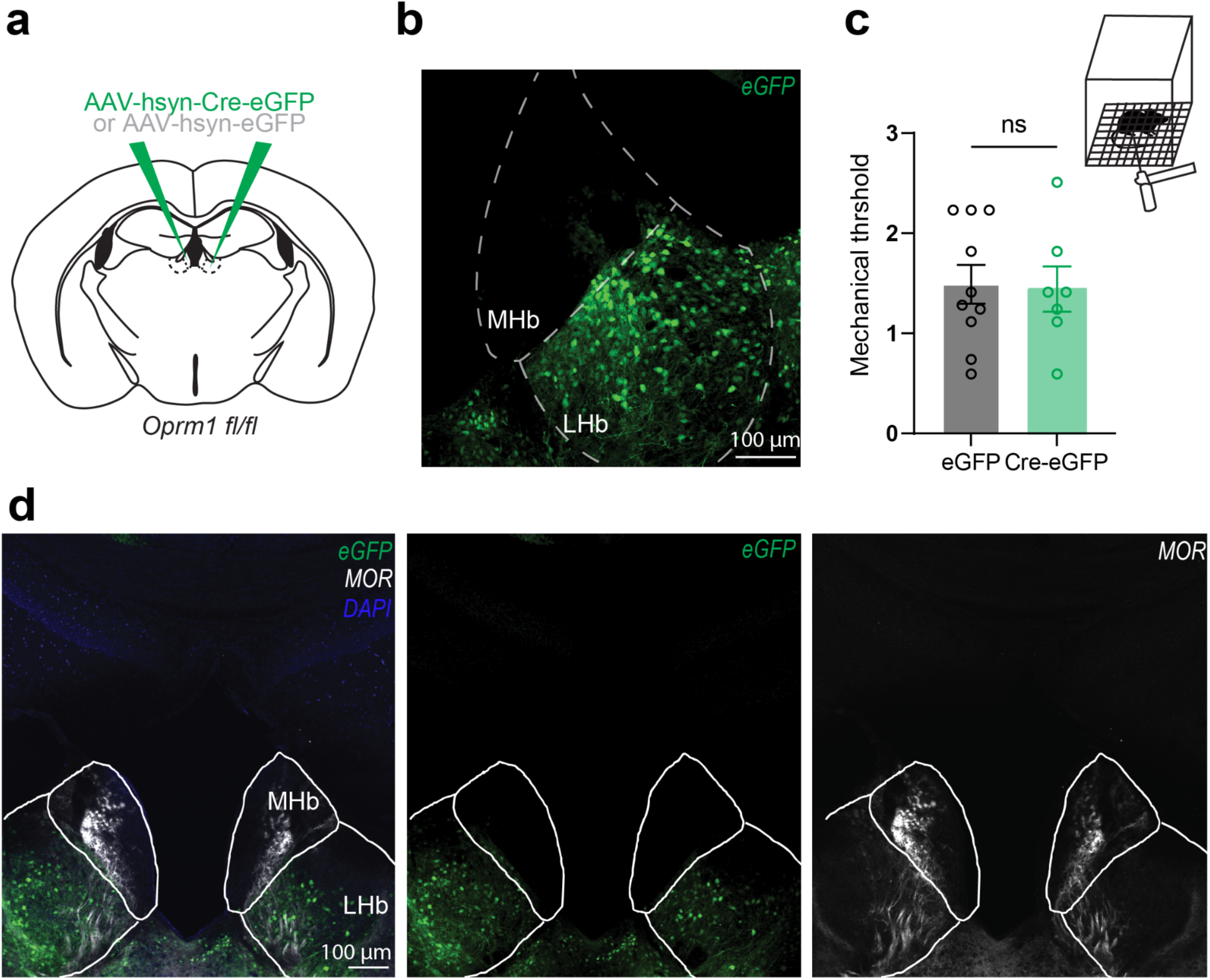
Minimal eGFP reporter expression in the MHb using hSyn-driven approach. (a) Schematic representation of viral injection sites in Oprm1^fl/fl^ mice, using either Cre-expressing or control viruses. (b) Representative coronal brain section from a control mouse showing eGFP expression. (c) Mechanical Von Frey threshold (unpaired, two-tailed, t-test, p = 0.8796, n=10 eGFP, 7 Cre). (d) Representative coronal brain section from a control mouse, showing MOR immunoreactivity and reporter expression.

**Supplementary Figure 3.**
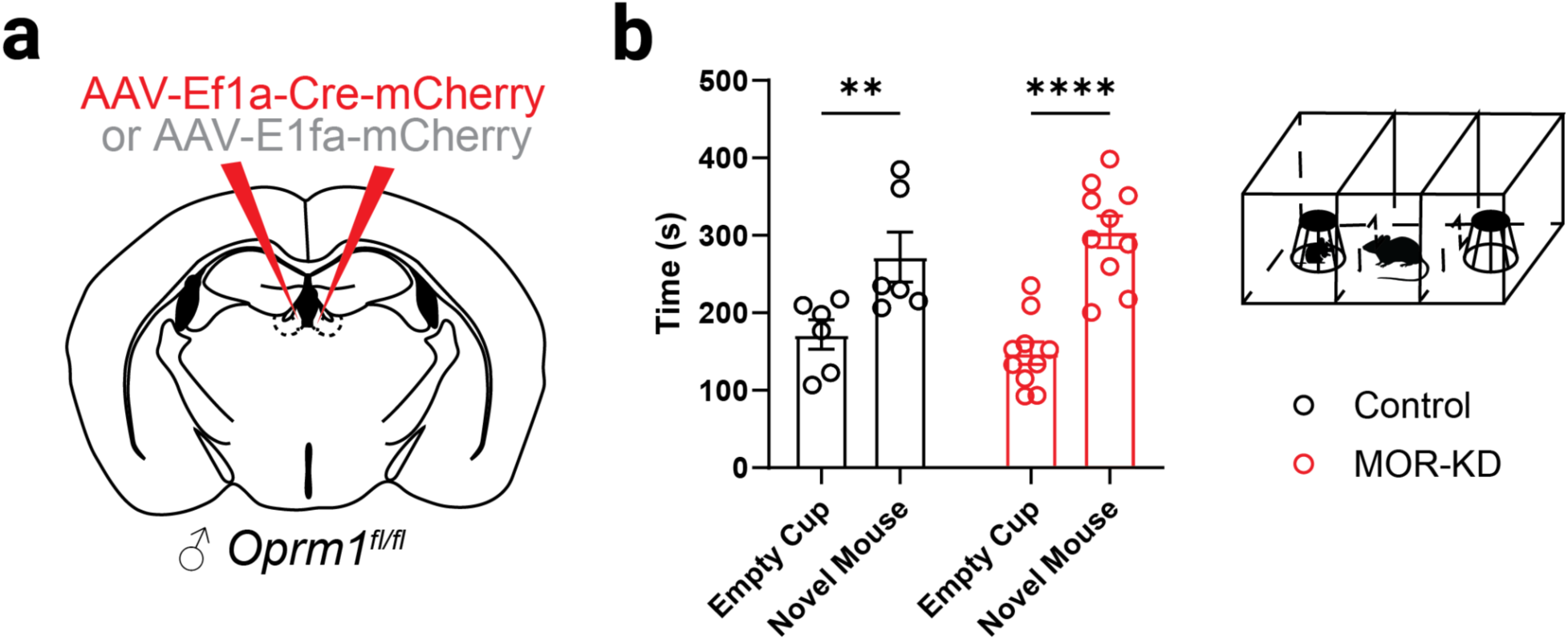
Targeted knockout of *Oprm1* in the habenula does not lead to decreased sociability. (a) Schematic representation of viral injection sites in Oprm1^fl/fl^ mice using control virus. (b) Social interaction total time (two-way ANOVA, F (1, 28) = 1.696, p=0.2034 for interaction of knockout and social stimulus, F (1, 28) = 34.97, ****p<0.0001 for main effect of social stimulus, F (1, 30) = 0.038, p=0.03824 for main effect of knockout, n=6 Control, 12 MOR-KO).

